# Examining Blunted Initial Response to Reward and Recent Suicidal Ideation in Children and Adolescents Using Event-Related Potentials: Failure to Conceptually Replicate Across Two Independent Samples

**DOI:** 10.1101/2020.05.19.104208

**Authors:** Austin J. Gallyer, Kreshnik Burani, Elizabeth M. Mulligan, Nicholas Santopetro, Sean P. Dougherty, Min Eun Jeon, Brady D. Nelson, Thomas E. Joiner, Greg Hajcak

## Abstract

A recent study by Tsypes and colleagues (2019) found that children with recent suicidal ideation had blunted neural reward processing, as measured by the reward positivity (RewP), compared to matched controls, and that this difference was driven by reduced neural responses to monetary loss, rather than to reward. Here, we aimed to conceptually replicate and extend these findings in two samples (*n* = 264, 27 with suicidal ideation; and *n* = 314, 49 with suicidal ideation at baseline) of children and adolescents (11 to 15 years and 8 to 15 years, respectively). Results from both samples showed no evidence that children and adolescents with suicidal ideation have abnormal reward or loss processing, nor that reward processing predicts suicidal ideation two years later. The results highlight the need for greater statistical power, as well as continued research examining the neural underpinnings of suicidal thoughts and behaviors.

---

Suicide is a growing problem among children and adolescents. In the United States from 1999 to 2018, the suicide rate increased from 1.2 to 2.9, and from 8.0 to 11.4, per 100,000 for those 10–14 and 15–19 years old, respectively, primarily among females (Centers for Disease Control, 2018). Moreover, suicide is the second leading cause of death for children and adolescents between 10 and 19 years of age (Centers for Disease Control and Prevention, 2018). Though death by suicide is understandably the focus of suicide prevention efforts, suicidal ideation is itself a serious public health issue among children and adolescents and one deserving of inquiry (Jobes & Joiner, 2019). The necessity of addressing suicidal ideation itself is underscored by the prevalence of suicidal ideation in this age range, with a recent survey of high school students in the United States finding that 18.8% reported seriously considering suicide during the past year (Ivey-Stephenson et al., 2020). Unfortunately, recent meta-analytic work shows that, while treatments for suicidal ideation do have an effect, effect sizes are small and are least effective in younger patients, such as children and adolescents (*g* = −.09; Fox et al., 2020). It has been suggested that this relative lack of treatment effectiveness may be due to failures to determine the causes of suicidal thoughts and behaviors and an overemphasis on risk factors that have modest longitudinal associations with suicidal ideation and other suicide-related outcomes (Fox et al., 2020; Franklin et al., 2017, 2018).

One approach that has been offered as a solution to developing more effective treatments for suicidal ideation is to study its purported causes from a transdiagnostic perspective grounded in neuroscience (Glenn et al., 2017). In particular, the Research Domain Criteria (RDoC), a framework developed by the National Institute of Mental Health that aims to integrate multiple levels of analysis to identify the underlying neural mechanisms of psychopathology, has been proposed as a way to improve the understanding of suicidal ideation (Glenn et al., 2017; Insel et al., 2010). One of the systems outlined by RDoC that has been suggested to be important for understanding suicidal ideation is the positive valence system. Specifically, differences in brain areas involved in reward processing have been proposed as important for the development of suicide-related outcomes (Auerbach et al., 2021; Schmaal et al., 2019), including suicidal ideation (Albanese & Hajcak, 2021). One strong, indirect source of support for this hypothesis comes from a meta-analysis that found that anhedonia is positively associated with suicidal ideation, independent of depression (Ducasse et al., 2017). However, the application of this finding in the treatment of suicidal ideation in children and adolescents requires evidence demonstrating differences in the reward system are associated with suicidal ideation in children and adolescents. Consequently, doing so requires an understanding of the brain structures involved in the reward system and the reward system’s development in late childhood and early adolescence.

## The reward system and its development in children and adolescents

The mesolimbic dopamine pathway plays a key role in the reward system (Berridge & Robinson, 1998) and consists of structures such as the ventral tegmental area, nucleus accumbens, and the amygdala. The reward system also includes cortical areas, such as the anterior cingulate cortex and the orbitofrontal cortex, among others (Haber & Knutson, 2010). Though what exactly this circuit is signaling about reward is debated (particularly the dopamine signal in the ventral tegmental area and ventral striatum; see Berridge, 2012; Schultz, 2016), it is scientific consensus that this circuit does not primarily mediate hedonic “liking” or pleasure (Berridge & Robinson, 1998, 2003). Research has converged on the theory that, beginning in late childhood and into early adolescence, the reward system undergoes significant changes that lead to increases in reward sensitivity (Galvan, 2010). For example, in rats, dopamine levels in the ventral striatum increase into adolescence, with levels dropping off in adulthood (Teicher et al., 1993). Dopamine receptor binding has also been shown to peak in adolescent rodents (Tarazi et al., 1998, 1999). Functional magnetic resonance imaging (fMRI) studies have also found that, compared to children (aged 7–12) and adults (aged 18–29), adolescents show hyperresponsive ventral striatal activity to receiving rewards (Galvan et al., 2006; Van Leijenhorst et al., 2010). There is also evidence from work using event-related potentials (ERPs) that reward sensitivity increases from late childhood into early adolescence (Burani et al., 2019).

Briefly, ERPs are direct measures of the neural response to specific events (e.g., onset of feedback) recorded using the electroencephalogram. The reward positivity (RewP) is an ERP that indexes the neural response to reward receipt, typically measured as the difference between monetary rewards and losses (Proudfit, 2015). The RewP is related to self-report and behavioral measures of reward processing (Bress & Hajcak, 2013; Pizzagalli et al., 2005) and is correlated with activation in reward-related brain regions, including the ventral striatum (Carlson et al., 2011). Recent work found that the RewP increases across early childhood into adolescence, with the youngest children seeing the greatest increases in the RewP at a two-year follow-up (Burani et al., 2019). A recent cross-sectional study that compared the RewP of children between 7-11 years old also found that older children had a larger RewP (Gibb et al., 2022). Together, these findings using animal models, fMRI, and the RewP, illustrate that late childhood and early adolescence are characterized by changes in the reward system leading to increased reward sensitivity, particularly in the ventral striatum. However, do differences in the reward system during this developmental period characterize children and adolescents who experience suicidal ideation?

## Reward system and suicidal thoughts and behaviors

A recent review claims that “blunted striatal activation characterize[s]” suicidal thoughts and behaviors (Auerbach et al., 2021). However, the evidence for this claim is relatively weak, particularly pertaining to whether this work has any bearing on suicidal ideation. For example, one study used a Cyberball peer-interaction task and found that adolescents with “high suicidal ideation” had blunted activity in the putamen, rather than the ventral striatum, compared to adolescents with “low suicidal ideation” to social rewards (Harms et al., 2019). Of note, adolescents in the “high suicidal ideation” group included those who had previously attempted suicide, limiting the applicability of the study’s findings to suicidal ideation research. Theoretically and empirically, risk factors for suicidal ideation and those for transitioning to suicidal behavior do not necessarily overlap (Joiner, 2005; May & Klonsky, 2016; O’Connor, 2011; Van Orden et al., 2010). Thus, it is possible that the blunting of the putamen was a result of adolescents who had previously attempted suicide, rather than a result of suicidal ideation per se. Similarly, another study found blunted striatal activity during a monetary incentive delay task in a group that had many adolescents who reported suicidal ideation, compared to a control group; however, the inclusion criteria for this group included engagement in non-suicidal self-injury (Sauder et al., 2016). While non-suicidal self-injury and suicidal thoughts and behaviors are related, factor analytic work has established that they are clearly distinguishable constructs with unique correlates (Evans & Simms, 2019), further complicating the distinction of the blunter striatal activity as a correlate of suicidal ideation as opposed to non-suicidal self-injury. Thus, these findings cannot address whether children/adolescents experiencing suicidal ideation have a blunted reward response compared to controls.

Work leveraging the RewP is also unclear. For example, one study found an *increased* RewP in depressed adolescents experiencing suicidal thoughts and behaviors compared to depressed controls (Pegg et al., 2020). However, in this study, suicidal thoughts and behaviors were conflated, again making it difficult to determine whether this difference in the RewP was driven by suicidal thoughts, suicidal behaviors, or both. Perhaps the best test of whether the reward system is blunted in children experiencing suicidal ideation was conducted by Tsypes and colleagues (2019). In this study, the authors found that children with recent suicidal ideation had a reduced RewP, compared to demographically and clinically matched controls (Tsypes et al., 2019). Moreover, the authors found that this difference was driven by an enhanced (i.e., more positive) neural response to losses. These results suggest that children with suicidal ideation have blunted reward processing, compared to children without suicidal ideation, implying that RewP differences may help identify, and potentially cause, children who develop suicidal ideation. These results also point to neural responses to losses as a potential marker of suicidal ideation among children. If the findings by Tsypes and colleagues (2019) can be replicated in multiple samples, including the effect size found (i.e., Hedges’ *g* = 0.60), it would provide strong evidence that the RewP has a unique relationship to suicidal ideation, given that previous findings in samples that conflated suicidal ideation and self-injurious behaviors did not find a similar diminishing of the reward response (e.g., Pegg et al., 2020). As noted above, suicidal ideation is prevalent in the general population, and is itself a state that can cause significant distress and impairment, even if an individual never engages in suicidal behavior (Jobes & Joiner, 2019).

Given the significance and implication of Tsypes and colleagues’ (2019) findings to the field’s understanding of suicidal ideation in late childhood, we sought to conceptually replicate and extend their findings. Specifically, we examined whether the RewP was blunted in children and adolescents with recent suicidal ideation, compared to controls, using similar identical methods, measures, and processing procedures as those used by Tsypes and colleagues in two large samples. Moreover, we sought to expand on Tsypes and colleagues’ (2019) findings by examining whether the RewP is not only related to suicidal ideation cross-sectionally but also whether the RewP could predict suicidal ideation two years later. In our first sample, consistent with Tsypes et al. (2019), we hypothesized that children and adolescents with recent suicidal ideation would have a significantly smaller RewP than those without recent suicidal ideation and that this effect would be driven by a blunted neural response to loss trials. For our second sample, we hypothesized the following: (1) Children and adolescents with recent suicidal ideation would have a significantly smaller RewP than those without recent suicidal ideation cross-sectionally at both Wave 1 and at Wave 2; and (2) RewP at Wave 1 would be negatively associated with suicidal ideation at Wave 2, two years later.

## Study 1: Conceptual Replication

### Methods

#### Open Science Statement

None of the analyses or data collected were pre-registered. All code used for our primary analyses, power analyses, supplementary materials, and figures are available at: https://osf.io/j6u2d/.

#### Participants

Our first sample consisted of 275 children and adolescents, but only 264 had complete data needed for analyses. Thus, the final sample size for Study 1 in all analyses was *n* = 264. Participants ranged from 11.01 to 14.98 years of age (*M* = 12.91, *SD* = 1.14). Our sample consisted of slightly more males than females (54.9% male). In this sample, 72.36% identified as white, 10.18% as African American, 6.18% as Hispanic, 4.73% as Asian, and 6.55% identified their race/ethnicity as “other.” Notably, our sample’s racial and gender demographics were similar to Tsypes and colleagues’ (2019) sample, but our sample was slightly older. Based on the present study’s sample size and an alpha of .05, we were powered at .84 to detect an effect size of Hedges’ *g* = 0.60 (calculation based on information in Tsypes et al. [2019]) for our difference in the RewP between those with recent suicidal ideation and controls. Participants and their parents provided informed consent and assent. The study was approved by the Institutional Review Broad at Florida State University.

#### Measures

##### Suicidal Ideation Measure

To assess for the presence of recent suicidal ideation (i.e., within the past two weeks), we used item 9^1^ of the original Children’s Depression Inventory (CDI; Kovacs, 1981) and the Schedule for School-Age Children—Present and Lifetime Version (K-SADS-PL; Kaufman et al., 1997). Individuals who endorsed a “1” (*I think about killing myself but I would not do it*) or “2” (*I want to kill myself*) on the CDI were grouped as having recent suicidal ideation. If individuals answered “1” or “2” on the CDI, then they were assessed for suicidal thoughts within the last two weeks on the K-SADS-PL interview, during which the children and their parents were asked, “Sometimes children who get upset or feel bad think about dying or even killing themselves. Have you/your child ever had these types of thoughts?” All children who endorsed suicidal ideation on the CDI also endorsed suicidal ideation during the K-SADS-PL. This is a small but important difference between this sample and the study by Tsypes and colleagues (2019), who grouped children as having recent suicidal ideation if they endorsed it on the CDI *or* the K-SADS-PL. Using our approach, 27 (10.23%) participants endorsed having recent suicidal ideation in this sample. For a breakdown of demographics between those with and without recent suicidal ideation, see Table 1.

**Table 1.** Demographics of Groups

| Study 1 |  |  |
| --- | --- | --- |
| Variable | No SI ($n = 237$) | Recent SI ($n = 27$) |
| Age | $M = 12.86, SD = 1.14$ | $M = 13.27, SD = 1.17$ |
| Sex (% female) | 44.30% | 51.85% |
| Race (% White) | 74.26% | 59.26% |
| Household Income (Median) | \$85,250 | \$80,500 |
| Study 2 Wave 1 |  |  |
| Variable | No SI ($n = 265$) | Recent SI ($n = 49$) |
| Age | $M = 12.29, SD = 1.80$ | $M = 13.09, SD = 1.60$ |
| Sex (% female) | 100% | 100% |
| Race (% White) | 85.17% | 81.25% |
| Household Income (Median) | \$105,000 | \$127,500 |
| Study 2 Wave 2 |  |  |
| Variable | No SI ($n = 222$) | Recent SI ($n = 33$) |
| Age | $M = 14.21, SD = 1.85$ | $M = 15.51, SD = 1.20$ |
| Sex (% female) | 100% | 100% |
| Race (% White) | 83.78% | 78.79% |
| Household Income (Median) | \$125,000 | \$90,000 |

*Note.* SI = suicidal ideation.

##### Doors Task

To measure neural responses to reward, we used the doors task, which is the same monetary guessing task used by Tsypes and colleagues (2019). The doors task is frequently used in ERP studies of reward processing (Bress et al., 2012, 2013; Burani et al., 2019; Klawohn et al., 2020). During this task, participants are presented with two doors and instructed to guess which door has the monetary prize. To make their selection, the participants use left and right mouse buttons corresponding with the door on the left and right, respectively. After selecting a door, visual feedback is given consisting of either a green arrow pointing upward, indicating that the participant won $0.50, or a red arrow pointing downward, indicating that the participant lost $0.25. Feedback was presented for 2,000 ms; then, the message “Click for next round” was presented. Our task consisted of 60 trials presented in two blocks of 30 trials. For this task, 30 win and 30 loss trials were presented in random order.

##### ERP Data Collection and Processing

During the doors task, electroencephalogram (EEG) was recorded with an active electrode EEG-system (ActiCHamp, Brain Products GmbH) with 32 electrodes positioned in accordance with the 10/20-system (ActiCAP, Brain Products GmbH). Electrode Cz served as the online reference, a ground electrode was placed on the forehead, and two electrodes were placed on the mastoids. Electrooculogram was recorded from four additional electrodes: two 1 cm above and below the left eye and two at the outer canthi of both eye. Continuous EEG waves were recorded at a sampling rate of 1000 Hz using a bandpass recording filter of 0.01–100 Hz. EEG data were processed using Brain Vision Analyzer, Version 2.1 (Brain Products, Gilching, Germany). For some participants (*n =* 14), at least one channel had to be interpolated using surrounding channels. Then, for all participants, data were re-referenced to the average of the mastoid electrodes and filtered from 0.01 to 30 Hz (Butterworth, 4^th^ order). We then extracted 1000-ms feedback-locked epochs, starting 200 ms before feedback presentation. Next, we corrected for eye movement artifacts using the algorithm developed by Gratton, Coles, and Donchin (1983). After this correction, segments that contained voltage steps > 50 mV between sample points, a voltage difference of 175 mV within a 400-ms interval, or a maximum voltage difference of < 0.5 mV within 100-ms intervals were automatically rejected.

Win and loss trials were averaged separately, and a baseline correction was applied using the 200-ms preceding feedback onset. After preprocessing, the mean, standard deviation, and range of usable trials for gain and loss were as follows: gain segments mean = 29.99 (SD = 0.09); 29 to 30; loss segments mean = 29.99 (SD = 0.18); 27 to 30. After processing, in line with Tsypes and colleagues’ (2019) procedures, we exported these averages for temporospatial principal component analysis (PCA), a factor analytic approach used to parse the ERP waveform into underlying constituent components. PCA was conducted using the ERP PCA Toolkit (Version 2.82; Dien, 2010a). In line with evidence-based recommendations (Dien, 2010b), we first conducted a temporal PCA using promax rotation. Covariance matrix and Kaiser normalization were used for this PCA (Dien et al., 2005). Based on the resulting scree plot, 15 factors were extracted in the temporal domain. We then analyzed the spatial distributions of these temporal factors by conducting a spatial PCA using infomax rotation (Dien, 2010b). Based on the averaged scree plot for all 15 temporal factors, we extracted four spatial factors, yielding 60 factor combinations. Twenty factors each accounted for more than 0.50% of the variance and were retained for further inspection (Kaiser, 1960). After examining the factors, TF03SF1 (i.e., temporal factor 3, spatial factor 1), which accounted for 11.20% of the variance, most closely resembled the RewP: this factor was maximal at the Cz electrode site at 235 ms following feedback onset and was more positive following win than loss trials (see Figure S2).

##### Data Analytic Plan

All statistical analyses were conducted in R (Version 3.5.2; R Core Team, 2018), using the following packages: “tidyverse” (Version 1.3.0; Wickham et al., 2019), “haven” (Version 2.2.0; Wickham & Miller, 2019), “afex” (Version 0.23-0; Singmann et al., 2019), “here” (Version 0.1; Müller, 2017), “DataExplorer” (Version 0.7.0; Cui, 2018), “emmeans” (Version 1.3.2; Lenth, 2019), and “psych” (Version 1.8.4; Revelle, 2018). To test our hypotheses, we conducted a 2×2 (condition: win, loss; ideation: no SI, recent SI) mixed-measures ANOVA^2^. To rule out alternative explanations for our results, we reran this ANOVA adding age as a covariate. We also reran this ANOVA adding household income as a covariate. Though we view our relatively larger sample size as a strength of our study, similar to Tsypes and colleagues’ (2019) study, we also reran our 2×2 mixed-measures ANOVA after selecting controls using a matching procedure. Specifically, we used a genetic matching algorithm to select two controls for every case with recent suicidal ideation using the “Matching” package (Version 4.9-7; Diamond & Sekhon, 2013; Sekhon, 2011). Similar to Tsypes and colleagues (2019), we also matched by age, sex, race, household income, lifetime history of major depressive disorder, lifetime history of any anxiety disorder, and current symptoms of depression, anxiety, and externalizing disorders. In the supplementary material, we also split the sample by gender and conducted our primary analyses. Last, given the results of our primary analyses and that traditional null hypothesis testing cannot provide confirmatory evidence for the null hypothesis, we decided to conduct two equivalence tests using the “TOSTER” package (Version 0.3.4; Lakens, 2017). These analyses require specifying an effect size range that we deem to be uninteresting or “effectively zero.” A significant equivalence test provides evidence that the effect is smaller than the specified effect size range. We chose two effect size ranges, resulting in two separate equivalence tests: (1) lower bound *d* = −0.24 and upper bound *d* = 0.24 and (2) lower bound *d* = −0.60 and upper bound *d* = 0.60. Our first effect size estimate was based on recent meta-analytic evidence that the upper bound on the relationship between ERPs and suicidal ideation is Hedges’ *g* = 0.24, and the second effect size range was based on the effect reported in the study we sought to conceptually replicate (Gallyer et al., 2021; Tsypes et al., 2019). For our equivalence tests, we used the PCAΔRewP as our dependent variable and ideation group (no SI, recent SI) as the independent variable.

## Results

The means, standard deviations, and sample sizes of study variables across Study 1 and Study 2 are presented in Table 2. The mixed-measures ANOVA revealed an independent effect of condition (*F*[1, 262] = 45.61, *p* < .001, η = .024), such that wins (*M* = 17.9, 95% CI [16.80, 19.00]) were greater than losses (*M* = 13.80, 95% CI [12.80, 14.9]). There was no independent effect of ideation (*F*[1, 262] = 1.66, *p* = .200, η = .005) or interaction between ideation and condition (*F*[1, 262] = 0.08, *p* = .773, η < .001; see Figure 1). Our ANCOVA with age as a covariate revealed no statistically significant effects, including the interaction between ideation and condition (*F*[1, 260] = 0.28, *p* = .601, η < .001). We also conducted our main ANOVA and ANCOVA with age as a covariate in male-identifying and female-identifying people separately (Supplement 1). Our findings were similar in both genders. Our analysis adding household income as a covariate revealed a significant difference between wins and losses (*F*[1, 232] = 6.87, *p* = .009, η = .004) but no other interactions or main effects. Our ANOVA using our matched controls based on the genetic matching algorithm also did not find a main effect of ideation (*F*[1, 64] = 2.49, *p* = .120, η = 0.033) or an interaction between ideation and condition (*F*[1, 64] = 0.25, *p* = .617, η < 0.001). The result of our first equivalence test, using effect size bounds of *d* = −0.24 and *d* = 0.24, was not significant (*t*[34.47] = −0.93, *p* = .179). In contrast, our second equivalence test, using effect size bounds of *d* = −0.60 and *d* = 0.60, was significant (*t*[34.47] = −2.82, *p* = .004), suggesting that, if there is an effect, it is smaller than that previously reported by Tsypes and colleagues (2019).

**Table 2.** Means and Standard Deviations of Study Variables

| Study 1 |  |  |
| --- | --- | --- |
|  | *M(SD)* | *M(SD)* |
| Variable | No SI (*n* = 237) | Recent SI (*n* = 27) |
| Wins | 18.20(8.18) | 16.09(7.91) |
| Losses | 13.95(7.91) | 12.18(7.44) |
| ΔRewP | 4.26(6.03) | 3.91(5.20) |
| Study 2 Wave 1 |  |  |
|  | *M(SD)* | *M(SD)* |
| Variable | No SI (*n* = 265) | Recent SI (*n* = 49) |
| Wins | 3.81(6.45) | 3.41(5.12) |
| Losses | -0.21(6.03) | -1.58(5.40) |
| ΔRewP | 4.02(5.99) | 4.99(5.14) |
| Study 2 Wave 2 |  |  |
|  | *M(SD)* | *M(SD)* |
| Variable | No SI (*n* = 222) | Recent SI (*n* = 33) |
| Wins | 6.64(6.14) | 6.55(4.52) |
| Losses | 2.70(5.70) | 2.20(4.43) |
| ΔRewP | 3.94(6.75) | 4.36(5.32) |

*Note.* SI = suicidal ideation; wins, losses, and ΔRewP all refer to the reward positivity (RewP), an event-related potential that indexes neural responses to reward receipt.

**Figure 1.**
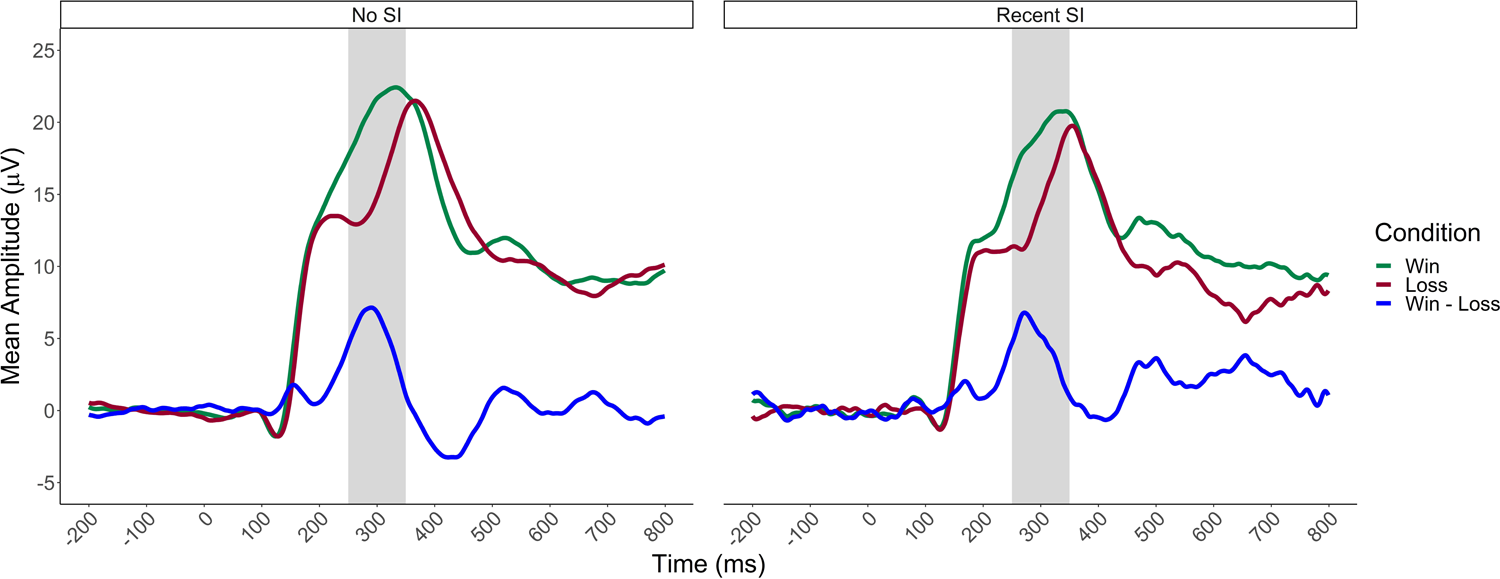
*Study 1 Stimulus-Locked Event-Related Potentials*, *Note.* Waveforms represent event-related potentials to wins and losses at Cz without conducting temporosptatial PCA. SI = suicidal ideation.

**Figure 2.**
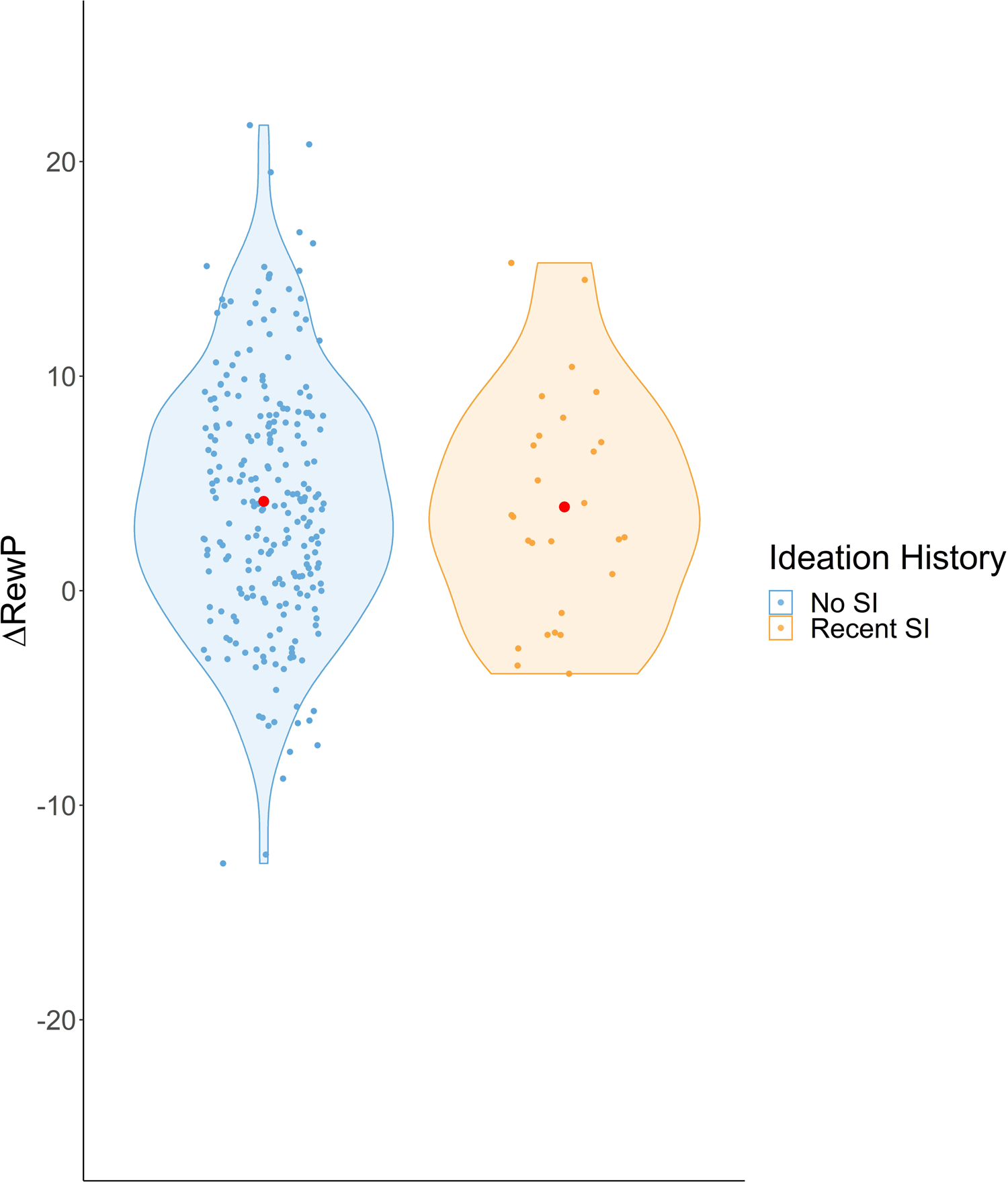
*Study 1* Δ*RewP of Children and Adolescents With and Without Recent SI*, *Note.* Red point represents mean RewP in each group. RewP = reward positivity, an event-related potential that indexes neu^Δ^ral responses to reward receipt; SI = suicidal ideation.

**Figure 3.**
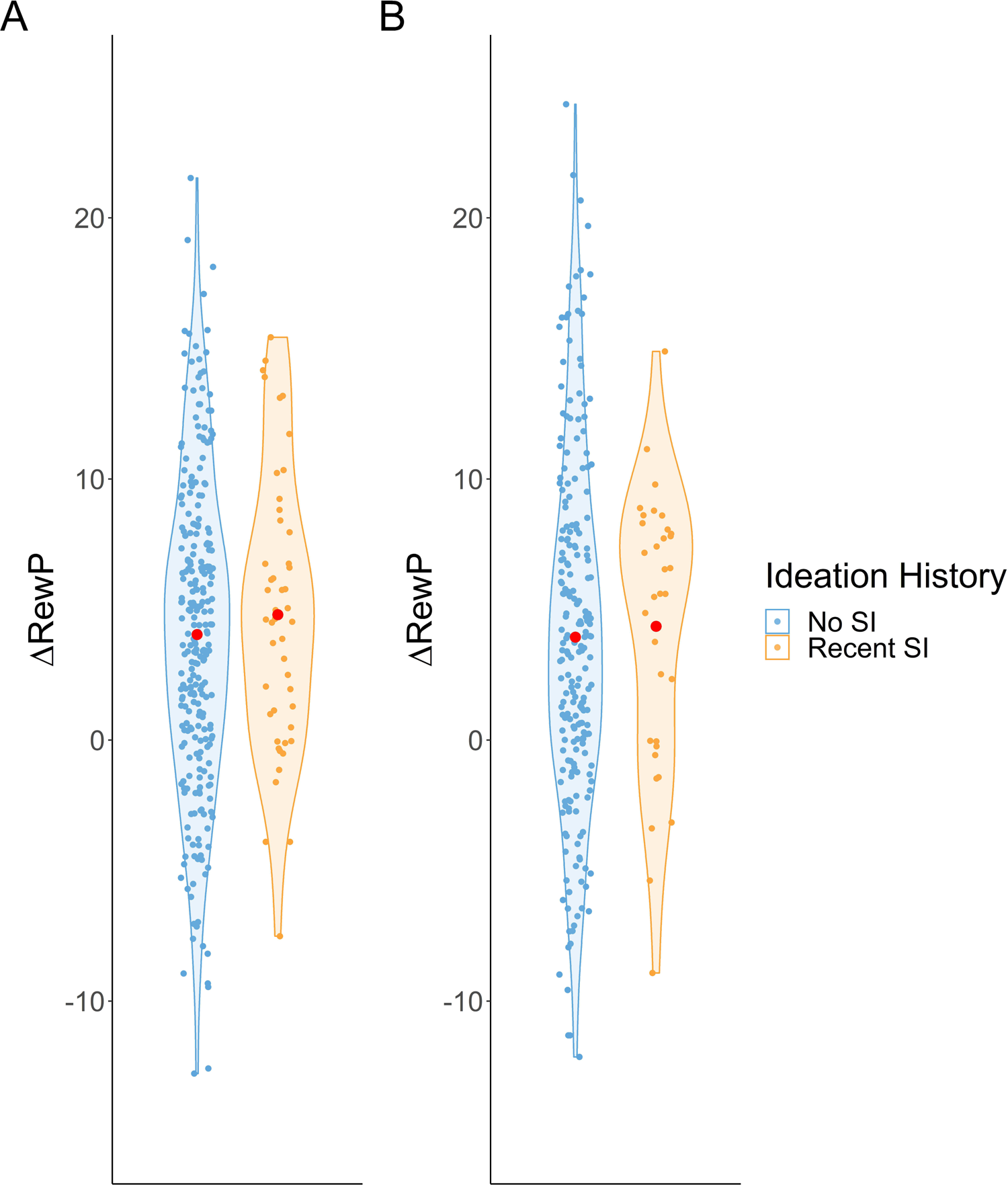
*Study 2* Δ*RewP of Children and Adolescents Across Wave 1 and Wave 2*, *Note*. Red points represent mean RewP across groups. A = Wave 1; B = Wave 2. ΔRewP = reward positivity, an event-related^Δ^potential that indexes neural responses to reward receipt; SI = suicidal ideation.

## Study 2: Extension

### Method

#### Open Science Statement

None of the analyses or data collected were pre-registered. All code used for our primary analyses, power analyses, supplementary materials, and figures are available at: https://osf.io/j6u2d/.

#### Participants

Our second sample consisted of 328 child and adolescent females, but only 314 had complete data needed for analyses. Therefore, all analyses for Wave 1 had a sample size of *n* = 314. These participants ranged from 8.01 to 15.04 years of age (*M* = 12.41, *SD* = 1.79) at Wave 1. At the two-year follow-up, 267 people returned to participate, but only 255 had complete data needed for analyses. Thus, the sample size at Wave 2 consisted of *n* = 255. At Wave 2, the participants age ranged from 9.89 to 17.18 years of age (*M* = 14.38, *SD* = 1.83). The original purpose of this sample was to study adolescent girl’s pubertal development and whether it was associated with depression. The analyses in this manuscript are unique to all previous use of this data. Notably, there were no differences in race (*Χ*^2^[12] = 4.85, *p* = .963) between those who dropped out of the study and those who returned at Wave 2 for the follow-up, but those who dropped out of the study were on average 0.71 years older than those who remained in the study (*t*[129.77] = 3.09, *p* = .013). In this sample, 81.54% identified their race as White/Caucasian, 5.85% as Black/African American, 0.31% as Native Hawaiian/Pacific Islander, 0.31% as American Indian/Alaskan Native, and 6.15% identified their race as “Other.” As in Study 1, this racial breakdown is similar to the sample in Tsypes and colleagues (2019), and our sample was slightly older than the participants recruited by Tsypes and colleagues. Participants were recruited from the community through a commercial mailing list, using flyers, and using word-of-mouth. Based on the present study’s sample size and an alpha of .05, we were powered at .97 to detect an effect size of Hedges’ *g* = .60 (calculation based on information in Tsypes et al. [2019]) for a difference in the RewP between those with recent suicidal ideation and controls at Wave 1, and .90 at Wave 2. All participants and their parents provided informed consent and assent, as approved by the Institutional Review Board at Stony Brook University.

#### Measures

##### Suicidal Ideation Measure

We used item 9 of the CDI and the K-SADS-PL interview to assess recent suicidal ideation, and participants who endorsed above “0” and/or endorsed suicidal ideation during the K-SADS-PL were categorized as having recent suicidal ideation. In contrast to Study 1, this coding is in line with Tsypes and colleagues’ (2019) approach. Based on this approach, 49 participants (15.6%) endorsed recent suicidal ideation at Wave 1, and 33 participants (12.9%) endorsed recent suicidal ideation at Wave 2.

##### Doors Task

We used the same doors task that was used in Study 1.

##### ERP Data Collection and Processing

During the doors task, continuous EEG was recorded using the ActiveTwo BioSemi system (BioSemi, Amsterdam, Netherlands) with a cap containing 34 electrodes placed according to the 10/20 system (i.e., 32 channels plus FCz and Iz). We also placed electrodes above and below the left eye, as well as near the outer canthi of both eyes, to measure vertical and horizontal electrooculographic activity, respectively. Two electrodes were also placed on the left and right mastoids. The EEG signal was preamplified at the electrode, and data were digitized at a 24-bit resolution with a sampling rate of 1024 Hz using a low-pass fifth-order sinc filter with a half-power cutoff of 204 Hz. Active electrodes were measured online with reference to a Common Mode Sense active electrode constructing a monopolar channel.

Data were processed using BrainVision Analyzer, Version 2.1 (Brain Products, Gliching, Germany). The data processing procedures were largely the same as in Study 1, with small differences in artifact detection: Segments that contained voltage steps > 50 mV between sample points, a voltage difference of 300 mV within a segment, or a maximum voltage difference of < 0.5 mV within 100-ms intervals were automatically rejected. The mean, standard deviation, and range of usable trials were as follows: gain segments mean = 29.99 (SD = 0.16); 28 to 30; loss segments mean = 29.99 (SD = 0.15); 29 to 30. As in Study 1, we conducted temporospatial PCA using ERP PCA Toolkit (Dien, 2010a). To do so, we first conducted a temporal PCA with promax rotation. A covariance matrix and Kaiser normalization were used for this PCA (Dien et al., 2005). Based on the resulting scree plot, 22 temporal factors were extracted in the temporal domain. We then analyzed the spatial distributions of these temporal factors by conducting a spatial PCA using infomax rotation (Dien, 2010b). Based on the averaged scree plot for all 22 temporal factors, we extracted 4 spatial factors, yielding 88 factor combinations. Twenty-four factors each accounted for more than 0.50% of the variance and were retained for further inspection (Kaiser, 1960). TF07SF1^3^, which accounted for 0.8% of the variance, most closely resembled the RewP, as it was maximal at the FCz electrode site at 349 ms and more positive for win trials than for loss trials. Thus, factor scores from this PCA factor were used as the RewP measure for Wave 1 (see Figure S2).

We repeated this same process for Wave 2. The mean, standard deviation, and range of usable trials were as follows: gain segments mean = 30 (SD = 0.00); 30 to 30; loss segments mean = 30.00 (SD = 0.00); 30 to 30. Our temporal PCA resulted in 19 factors, and our spatial PCA resulted in three factors, yielding 57 factor combinations together. Eighteen factors each accounted for more than 0.50% of the variance and were retained for further inspection (Kaiser, 1960). The TF06SF1 factor, which accounted for 1.5% of the variance, most closely resembled the RewP, as it was maximal at the FCz electrode site at 278 ms and more positive for win trials than for loss trials (see Figure S2).

##### Data Analytic Plan

We used the same statistical software and packages that were used in Study 1. To test our hypotheses, we conducted the same analyses we used in Study 1 for Wave 1 and Wave 2 separately. One notable difference is that, for our genetic matching procedure, we did not match based on gender, given that the entire sample in this study was female. To extend on Tsypes and colleagues (2019) and other work (Pegg et al., 2020) that found a cross-sectional relationship between the RewP and suicidality in children and adolescents, we also conducted a series of logistic regression analyses. Specifically, we examined whether ΔRewP at Wave 1 predicted ideation status at Wave 2. Given work showing that childhood and adolescence are associated with significant development in reward processing, we also reran this analysis including age as a covariate (Burani et al., 2019). Last, to further investigate any interactions with gender, in the supplementary materials we combined Sample 1 and Sample 2 Wave 1 into a single random-effects analysis (see Table S1).

## Results

### Wave 1 Analyses

As expected, our 2×2 mixed-measures ANOVA revealed a main effect of condition (*F*[1, 309] = 95.56, *p* < .001, η^2^ = .067), such that wins (*M* = 3.91, 95% CI [3.16, 4.66]) were greater than losses (*M* = −0.59, 95% CI [-1.34, 0.16]). In line with our Study 1 results, we did not find evidence of a main effect of ideation (*F*[1, 309] = 1.11, *p* = .293, η^2^ = .003) or an interaction between ideation and condition (*F*[1, 309] = 1.11, *p* = .294, η^2^ = .001). Our ANCOVA with age as a covariate revealed a main effect of age (*F*[1, 307] = 7.36, *p* = .007, η^2^ = .018) but no other statistically significant effects, including the interaction between ideation and condition (*F*[1, 307] = 1.15, *p* = .285, η^2^ = .001). Our ANCOVA adding income as a covariate revealed the expected main effect of condition (*F*[1, 262] = 9.06, *p* = .003, η^2^ = .008), but no other effects were significant. Our analysis using our matched controls revealed a main effect of condition (*F*[1, 136] = 46.66, *p <* .001, η^2^ = .076), but our main effect of ideation (*F*[1, 136] = 0.78, *p* = .378, η = .004) and the interaction between ideation and condition (*F*[1, 136] = 0.67, *p* = .416, η = .001) were not significant. The results of our first equivalence test, using effect size bounds of *d* = −0.24 and *d* = 0.24, were not significant (*t*[72.37] = −0.93, *p* = 0.259). In contrast, our second equivalence test, using effect size bounds of *d* = −0.60 and *d* = 0.60, was significant (*t*[72.37 = 2.87, *p* = .003), again suggesting that, if there is an effect, it is smaller than previously reported by Tsypes and colleagues (2019).

### Wave 2 Analyses

Our 2×2 mixed-measures ANOVA revealed a main effect of condition (*F*[1, 253] = 45.59, *p* < .001, η = .056) such that wins (*M* = 6.70, 95% CI [5.87, 7.54]) were greater than losses (*M* = 2.56, 95% CI [1.72, 3.39]). In line with our Study 1 and Study 2 Wave 1 results, we did not find evidence of a main effect of ideation (*F*[1, 253] = 0.11, *p* = .739, η < .001) or an interaction between ideation and condition (*F*[1, 253] = 0.11, *p* = .736, η = .001). Our ANCOVA with age as a covariate revealed no statistically significant effects, including the interaction between ideation and condition (*F*[1, 251] < 0.01, *p* = .980, η < .001). Our ANCOVA adding income as a covariate revealed the expected main effect of condition (*F*[1, 228] = 9.06, *p* = .003, η = .013), but no other effects were significant. Our analysis using our matched controls revealed a main effect of condition (*F*[1, 91] = 20.79, *p <* .001, η = .056), but our main effect of ideation (*F*[1, 91] = 0.01, *p* = .927, η < .001) and the interaction between ideation and condition (*F*[1, 91] = 0.45, *p* = .505, η = .001) were not significant. The results of our first equivalence test, using effect size bounds of *d* = −0.24 and *d* = 0.24, were not significant (*t*[48.73] = 1.19, *p* = 0.120). In contrast, our second equivalence test, using effect size bounds of *d* = −0.60 and *d* = 0.60, was significant (*t*[48.73] = 3.14, *p* = .002), again suggesting that, if there is an effect, it is smaller than previously reported by Tsypes and colleagues (2019).

### Wave 1 ΔRewP Predicting Wave 2 Ideation

Our first logistic regression predicting Wave 2 ideation status using Wave 1 ΔRewP as the predictor was not significant (*OR* = 0.98, 95% CI [0.92, 1.04], *p* = 0.479). Our second logistic regression adding Wave 1 age as a covariate showed that Wave 1 age was a significant predictor of Wave 2 suicidal ideation status (*OR* = 1.62, 95% CI [1.26, 2.17], *p* < .001), but Wave 1 ΔRewP was still not significant (*OR* = 0.96, 95% CI [0.90, 1.02], *p* = 0.209).

## Discussion

The purpose of this study was to test whether children and adolescents with recent suicidal ideation have blunted neural response to gains and/or losses cross-sectionally in a large sample of males and females, and whether blunted reward processing predicts suicidal ideation two years later in a sample of females. A previous cross-sectional study found that children with suicidal ideation exhibit aberrant reward processing, compared to matched controls, and this difference appeared to be driven by neural responses to loss trials, rather than to gain trials (Tsypes et al., 2019). However, in our conceptual replication using quite similar methods, we did not find any evidence that children and adolescents with recent suicidal ideation had different neural reward responses than those without recent suicidal ideation. We attempted to rule out other explanations by conducting additional analyses, including controlling for age, controlling for household income, using matched samples, and by combining Study 1 and Study 2 Wave 1 into a single analysis (See Supplementary Material). Nevertheless, we were not able to find a main effect of suicidal ideation or an interaction between suicidal ideation group and wins/losses in a single analysis. To further investigate these null findings, we also conducted equivalence tests using two different cutoffs (viz., *d* = .60 and *d* = .24) and only found statistical equivalence when we used the larger cutoff. These results suggest that, *if* there is a difference in reward processing in children and adolescents within this age range, it is likely smaller than the effect originally reported by Tsypes and colleagues (2019).

Our results are in contrast to previous work that has found differences in striatal activation in adolescents with high suicidal ideation (Auerbach et al., 2021). However, aside from the fact that our study leveraged the RewP, rather than fMRI, there are some key methodological differences between our study and the broader literature that uses fMRI to study reward processing in suicidal thoughts and behaviors. First, many fMRI studies did not distinguish among suicidal ideation, suicidal behavior, and nonsuicidal self-injury. Moreover, many fMRI studies examine other types of rewards, such as social reward (e.g., Harms et al., 2019). Therefore, it is possible that these methodological differences explain the discrepancy between our results and the general fMRI literature. However, a recent, well-powered fMRI study (*N =* 4,074) failed to find any differences in any phase of reward processing (e.g., reward anticipation, reward receipt) in any brain region in children (9–10 years old) experiencing suicidal ideation, compared to controls (Vidal-Ribas et al., 2021). In fact, Vidal-Ribas and colleagues (2021) also used equivalence testing and found that these effects were statistically equivalent to their smallest effect size of interest (i.e., *d* = .15). Thus, our results are in line with one of the most well-powered fMRI studies to date on reward processing in children with suicidal ideation.

Regarding the study by Tsypes and colleagues (2019) and why we failed to find the same results that they did, we believe there are at least three possible explanations. First, there are some subtle but key demographic differences between our samples and the sample reported by Tsypes and colleagues. Namely, the average age of both of our samples was about two-to-three years older, and the median household income in our samples was much higher ($85,000–$125,000, depending on sample and wave of data collection), compared to the sample collected by Tsypes and colleagues (∼$20,000). The differences in age are especially key, as the reward system, and the RewP in particular, undergo substantial changes in late childhood and early adolescence, with reward sensitivity increasing over time (Burani et al., 2019; Galvan, 2010; Gibb et al., 2022; Teicher et al., 1993). Thus, it is possible that reward processing in children with suicidal ideation only differs from that of children without suicidal ideation in late childhood, and that this effect diminishes as children progress into adolescence. This assertion may be supported by a recent study that also failed to find a difference in the RewP in adults with a previous suicide attempt compared to controls (Tsypes et al., 2020). That is, given previous work has shown a difference in the RewP in children experiencing suicidal thoughts and behaviors (Pegg et al., 2020), these effects in both suicidal ideation and suicidal behaviors may only be present at an early age (however, it should be noted that Pegg et al., 2020 and Tsypes et al., 2019 found effects in the opposite direction). While we tried to rule this explanation out by conducting ANCOVA’s while controlling for age, this explanation still remains a possibility that will need to be examined in future research.

Moreover, given that the median incomes in our samples were much higher than that in Tsypes and colleagues’ (2019) sample, it is possible that environmental stressors such as poverty moderate the difference in reward processing between children and adolescents experiencing suicidal ideation. The potential for this type of effect is bolstered by research showing that poverty can influence the relationship between neural processing and other processes, such as inflammation (Miller et al., 2021).

A second explanation for the contrast in our findings is that differences in methodology may have contributed to the results, even despite our efforts to replicate those used by Tsypes and colleagues as closely as possible. Most notably, because Tsypes and colleagues’ study used PCA, it is difficult, if not impossible, to exactly replicate the weightings of time and space in the quantification of the RewP. Specifically, PCA may lead to construct invariance between studies, thereby decreasing their comparability. Though we chose the PCA factors that most closely resembled the RewP in each of our samples, our Study 1 PCA factor and Study 2 Wave 1 factor peaked at slightly different times than is traditionally attributed to the RewP (i.e., 235 ms and 349 ms vs. 250–350 ms). Therefore, it is possible that factor selection influenced our results. Nonetheless, we found null results across Study 1 and Study 2, even when we used area measures (250–350 ms) without conducting PCA. Further, though they did not formally analyze their data in this way, the figure of the non-PCA waveforms in Tsypes and colleagues (2019) appear to show a difference in the RewP between children with recent suicidal ideation and controls, whereas our non-PCA results also failed to find a difference in the RewP between the two groups. Therefore, while PCA non-invariance is a potential problem, this explanation is less likely to account for the discrepancy between our study and the study by Tsypes et al. (2019) if their non-PCA waveforms align with their PCA results. We also determined the groups slightly differently in Study 1: whereas Tsypes and colleagues (2019) determined children to have recent suicidal ideation if they endorsed suicidal ideation on the CDI *or* the K-SADS-PL, in Study 1 the suicidal ideation portion of the K-SADS-PL was only administered if participants first endorsed suicidal ideation on the CDI. In practice, this methodological difference means that it is possible we missed participants who would have endorsed suicidal ideation on the K-SADS-PL but not on the CDI; however, recent work among adults shows that this group is expected to be extremely small, given that increased anonymity is associated with a greater likelihood to endorse suicidal thoughts and behaviors (Deming et al., 2021). Given this evidence and the fact that we used the same approach to determine our recent suicidal ideation group as Tsypes and colleagues (2019) in our second sample and still did not find differences in the RewP between groups, it seems unlikely that this methodological difference drove our null results in our first sample.

The third potential explanation for the discrepancy between our findings and Tsypes and colleagues’ findings is that our study is a false negative (i.e., there is a true effect, but we failed to find it) or that the study by Tsypes and colleagues was a false positive. Only considering our study and the study by Tsypes and colleagues, it is difficult to decide between these two scenarios. We recently conducted a meta-analysis on the relationship between ERPs and suicidal ideation and found a very small (viz., Hedges’ *g* = −.06) blunting of the RewP in individuals experiencing suicidal ideation compared to controls (Gallyer et al., 2021). For reasons we discuss in that meta-analytic review, including the small effect size and the high correlation between depression and suicidal ideation (Rogers et al., 2016), it cannot be ruled out that this blunting of the RewP is a false positive. In isolation, this possibility would appear to suggest that there are no differences in reward processing between those with versus without suicidal ideation. However, it is still possible that our results, and the results of our meta-analysis, are due to insufficient sample sizes. Recent work has shown that, across both fMRI and ERP studies, sample sizes tend to be too small to detect realistic effect sizes in individual differences (Button et al., 2013; Clayson et al., 2019; Elliott et al., 2020; Nord et al., 2017)^4^.

What do our results mean about future work on reward processing in children and adolescents experiencing suicidal ideation? First, we recommend better measurement practices in the study of the neural correlates of suicidal thoughts and behaviors. As was used in our study, it is common practice to use single-item measures of suicidal thoughts and behaviors and to dichotomize variables into a group experiencing any suicidal thoughts and/or behaviors and another group experiencing no suicidal thoughts and/or behaviors. The use of a single-item assessment is particularly problematic given evidence that a non-negligible number of participants who endorse suicidal thoughts/behaviors on a single item do not meet standardized criteria for suicidal thoughts/behaviors if asked follow-up interview questions (Hom et al., 2016; Millner et al., 2015). Future work should ideally use validated multi-item interviews that allow for clarifying questions (e.g., Chu et al., 2015; Gallyer et al., 2020; Gratch et al., 2021; Joiner et al., 1999; Nock et al., 2007).

Second, we recommend that researchers carefully consider the ideation-to-action framework when studying suicidal thoughts and behaviors (Klonsky et al., 2018). A large portion of the fMRI and ERP literature was not directly comparable to our own study because many studies conflate suicidal thoughts, suicidal behaviors, and nonsuicidal self-injury (e.g., Harms et al., 2019; Pegg et al., 2020; Santamarina-Perez et al., 2019; Sauder et al., 2016). Distinguishing among suicidal ideation, suicidal behaviors, and nonsuicidal self-injury will aid in interpreting which neural correlates are, or are not, associated with different phases of the ideation-to-action framework.

Third, researchers may consider examining fMRI and ERPs in the same participants looking at different phases of the reward response. Such a study, if it were sufficiently powered, would provide an excellent test of whether reward processing plays a role in suicidal ideation in children and adolescents.

The current study has notable strengths. First, our study largely used the same methods as the study we attempted to conceptually replicate, including the same task, the same measure of recent suicidal ideation, and similar data processing and data analytic procedures. The present study also used relatively large samples that are not typically seen in ERP research of suicidal thoughts and behaviors, making our study well-powered to detect medium effects. Our study also used two independent samples and extended on previous work by using an additional longitudinal sample, with EEG and suicidal ideation measurements at both timepoints. As such, the current study is the first to our knowledge to attempt to prospectively predict suicidal ideation using an ERP—here, the RewP.

Despite these strengths, the current study also has some limitations. First, though our study was well-powered to detect the effect size previously found by Tsypes and colleagues, much larger samples are needed to detect smaller effects. Due to measurement error, it is expected that ERPs will have small-to-moderate effects with behavioral and self-report outcomes. Given the low base rate of suicidal ideation and other suicidal outcomes, large samples are crucial for elucidating the relationship between the RewP and suicidal thoughts and behaviors. Second, our second sample consisted only of female participants. Though this potential demographic limitation is mitigated by our first sample, which has a fairly even split of males and females, this fact underscores that our longitudinal analyses may not apply to samples that include males. Thus, examining whether our longitudinal results generalize to other genders, as well as studying any differences in response to reward and its relation to future suicidal ideation across genders, is an important area for future research.

## Conclusions

In summary, the present study attempted to conceptually replicate and extend previous findings that children with recent suicidal ideation had blunted neural reward processing, compared to controls. In two samples, we did not find evidence of this effect. We also did not find evidence that blunted reward processing was associated with future suicidal ideation. Our results highlight the need for larger sample sizes and multiple studies when investigating the relationship between neural reward processing and suicidal ideation. Our hope is that the present results will spur further research into the neural underpinnings of suicidal thoughts and behaviors and that greater collaboration and larger sample sizes will begin to dominate this field. Such approaches will greatly enhance our ability to contribute to the knowledge of the neurophysiology of suicidal thoughts and behaviors and will allow us to explore the potential clinical applications of event-related potentials to suicidal thoughts and behaviors.

## Author Contributions

Testing and data collection were performed by NS and KB. AJG performed the data analysis and interpretation under the supervision of TEJ and GH. AJG drafted the paper, and all other authors provided critical revisions. All authors approved the final version of the paper for submission.

## Footnotes

1 Note that we used the original CDI. In the revised CDI, the suicide item is item 8.

2 We also conducted these analyses using an area measure (250–350 ms) of the RewP at FCz, without using temporospatial PCA. The pattern of results remained the same.

3 Notably, another factor was also considered that explained more of the variance. We opted for this factor because: (1) The time window for the alternative factor was much earlier/outside of the traditional range of the RewP, and (2) the alternative factor did not display sufficient difference between wins and losses. Importantly, we also ran all of our analyses using the alternative factor; the results were the same: We found no differences in the RewP between the groups.

4 It is important to remember that low statistical power increases the probability of false negatives *and* false positives (see Button et al., 2013).

